# Genotype and environmental effects shape the house fly microbiome (*Musca domestica*)

**DOI:** 10.64898/2026.04.06.716741

**Authors:** Sohana Al Sanjee, Kiran Adhikari, Richard P. Meisel

## Abstract

Animal-associated bacteria (microbiomes) can have important effects on host phenotypes and fitness. Microbiomes can also vary across individuals in ways that depend on host genotype and environment. Temperature is an especially important environmental factor that can affect the microbiome in a way that depends on host genotype and affects organismal fitness. Thermal stress, in particular, can have dramatic effects on animal microbiomes, including dysbiosis and immune dysregulation. However, most previous work on extreme temperature effects has focused on heat stress. To investigate how low temperatures affect the microbiome of a warm-adapted animal, we characterized the bacterial communities associated with house fly (*Musca domestica*) males raised at cool (18°C) and warm (29°C) temperatures. We sampled two distinct genotypes in these experimental flies, each of which is associated with a particular thermal environment (warm or cool). We contrasted our experimental results with the microbiomes we characterized in wild house flies from two collection sites with different large animals present. We found that temperature has a much stronger effect on the house fly microbiome than the host genotype in our experimental flies. Consistent with the strong environmental effects in our experiment, we found that wild house fly microbiomes differed between the two collection sites. Despite these environmental effects on the house fly microbiome, we did not detect evidence for dysbiosis associated with either cool or warm temperatures. We therefore conclude that the environment has more of an effect on the house fly microbiome than host genotype, but dysbiosis does not occur within the temperature range we considered.

## Introduction

Animals harbor prokaryotic and eukaryotic microbes that affect ecologically and evolutionarily important organismal phenotypes (Lange et al., 2023; Lemoine et al., 2020). This host-associated microbiome impacts physiological processes, such as nutritional supplementation, tolerance to environmental perturbation, resistance to pathogens, and immune system priming (Bahrndorff et al., 2016; Cerf-Bensussan & Gaboriau-Routhiau, 2010). The effects of the microbiome on host phenotypes can also have important fitness-related consequences (Henry et al., 2024), which can affect local adaptation to specific environments (Walters et al., 2020) or shift selection pressures on the host (Rudman et al., 2019). In addition, specific bacterial genes have been identified that modulate the effect of the microbiome on the host, in a host genotype-specific manner (Chaston et al., 2014; Matthews et al., 2020).

Of particular interest are the relative contributions of environment and host genotype on animal-associated microbiomes (Benson et al., 2010; Goodrich et al., 2014; Qin et al., 2022; Scepanovic et al., 2019; Spor et al., 2011). For example, in *Drosophila melanogaster*, genotypes with adaptive and ecologically relevant phenotypic effects are predictive of differences in host-associated microbiomes (Kristensen et al., 2024; Wang et al., 2020), providing evidence that genotype may be more important than environment in shaping the microbiome. Consistent with that interpretation, the *D. melanogaster* microbiome is constant across an ecological cline, suggesting host filtering causes the microbiome to be robust to ecological effects (Henry & Ayroles, 2022). In contrast to *D. melanogaster*, there is no effect of *Aedes albopictus* genetic variation on the microbiome, but the environment may be important (Minard et al., 2015, 2018). These results suggest that the relative contributions of genotype and environment on animal microbiomes vary across host taxa.

If we consider the microbiome to be phenotype of the host organism, we can interpret microbiome differences within the context of genotype-by-environment (G×E) interactions, where the microbiome depends on the host genotype and the environmental context in which it is found (Via & Lande, 1985). For instance, among-population differences of the fire ant *Solenopsis invicta* gut microbiome are associated with host genotype and geographical distribution (Xiao et al., 2023). Similarly, in *Daphnia magna*, both genotype and temperature affect the microbiome, but differences among groups are not necessarily observed in the most abundant taxa (Frankel-Bricker et al., 2020; Sullam et al., 2018). The lack of consistent patterns in microbiome diversity across host taxa demonstrates that further work is needed to understand how host genotypes and environmental variation affect animal-associated microbiomes.

Temperature variation is an especially important environmental factor that affects animal microbiomes (Huus & Ley, 2021; Sepulveda & Moeller, 2020), especially for ectotherms such as insects (Mazzucco & Schlötterer, 2021; McMunn et al., 2022; Moghadam et al., 2018). As examples, temperature is one of the best predictors of the *D. melanogaster* microbiome sampled across the world (Gale et al., 2025), and water temperature greatly affects the microbiome of the sea anemone *Nematostella vectensis* (Baldassarre et al., 2023). The effects of temperature on animal microbiomes can also accompany changes in host physiology and life history traits, including survival (Arango et al., 2021). In the spring field cricket, *Gryllus veletis*, seasonal fluctuations in temperature are associated with changes in thermal tolerance, immune function, and the microbiome (Ferguson et al., 2018). The mode of action for these temperature-dependent effects likely depends on both host physiology and microbial function. In hosts, temperature can modulate immune activity, gut physiology, and metabolic processes, thereby altering the ecological niche available for microbial colonization (Moghadam et al., 2018). Similarly, microbial taxa differ in their thermal tolerance and functional responses, leading to shifts in community composition and activity under different thermal conditions (Sepulveda & Moeller, 2020).

Amongst all the temperature-dependent effects, animal microbiomes are particularly sensitive to thermal stress, and the associated changes to the microbiome can affect organismal traits (S. Li et al., 2022; Raza et al., 2020). For example, inter-strain differences in the damselfly *Ischnura elegans* associated bacterial community converge upon a similar microbiome with heat stress (Theys et al., 2023). In addition, high temperature stress affects the intestinal microbiome of the marine snail *Pomacea canaliculata*, while low temperature has a more moderate effect (S. Li et al., 2022). In *Drosophila subobscura*, heat stress decreases the diversity of the microbiome, and complete removal of the microbiome causes axenic flies to have lower heat tolerance (Jaramillo & Castañeda, 2021). While these results demonstrate how heat stress affects animal-associated microbiomes, it is not clear if microbiomes are disrupted by higher temperatures *per se* or more general deviation from the thermal optimum. Addressing that question requires shifting a warm adapted host to a cooler temperature and measuring the change in the associated microbial community.

To understand how the microbiome of a warm-adapted animal is affected by cooler temperatures in a genotype-specific manner, we compared the bacterial communities of house flies (*Musca domestica*) with two different genotypes raised at warm and cool temperatures. House fly is well-suited for this line of investigation because it is a warm-adapted animal with segregating genetic variation that affects temperature-dependent phenotypes (Adhikari et al., 2021; Delclos et al., 2021; Hamm et al., 2015). The house fly male-determining gene (*Mdmd*) can be found on at least four of the six chromosomes in natural populations (Sharma et al. 2017). The Y chromosome (Y^M^) and third chromosome (III^M^) are the most common male-determining chromosomes in natural populations (Hamm et al., 2015). Notably, their distributions follow a geographical cline—males from higher latitudes are more likely to carry Y^M^, and males from lower latitudes tend to have the III^M^ genotype (Hamm et al., 2005; Kozielska et al., 2008; Tomita & Wada, 1989). Consistent with this geographic distribution, males carrying the III^M^ chromosome have greater heat tolerance and prefer warmer temperatures, while Y^M^ confers cold tolerance and preference for cooler temperatures (Delclos et al., 2021).

House flies additionally have important ecological, developmental, and physiological interactions with bacteria throughout their life cycle (Nayduch & Burrus, 2017). House fly larvae live in close association with microbe-rich substrates, such as animal excrement and decomposing organic materials (West, 1951). The larvae also depend upon live microbes to complete their growth and development (Schmidtmann & Martin, 1992; Zurek et al., 2000). The microbial communities of house fly have been extensively sampled in both natural and laboratory environments (Bahrndorff et al., 2017; Junqueira et al., 2017; Monyama et al., 2023; Neupane, Park, et al., 2024; Voulgari-Kokota et al., 2022), and differences have been observed between sexes and across natural populations (Bahrndorff et al., 2020; Neupane et al., 2020; Neupane & Nayduch, 2022; Park et al., 2019). However, no prior work has directly tested for effects of temperature or house fly genotypes on fly-associated microbiomes.

We tested if genotype and temperature affect house fly microbiomes in laboratory flies, and we compared those differences to house flies sampled from natural populations. To those ends, we raised males carrying either the Y^M^ or III^M^ chromosomes at 18°C or 29°C, and we sequenced the bacterial 16S rRNA gene to quantify the bacterial taxa associated with the flies. In addition, we sampled house flies from two natural populations in Texas (USA), genotyped males for the Y^M^ and III^M^ chromosomes, and performed 16S rRNA sequencing to quantify microbiomes. Our results allowed us to evaluate the relative effects of host genotype and environment on the house fly associated microbiome.

## Materials and Methods

### House fly strains and rearing

We performed our experiment using two house fly strains, CSkab (III^M^ males) and IsoCS (Y^M^ males). Both strains share a common genetic background of the Cornell susceptible (CS) strain (Scott et al., 1996). IsoCS was previously created by crossing a Y^M^ chromosome from Maine onto the CS background (Hamm et al., 2009). CSkab was created by backcrossing the III^M^ chromosome from the KS853 strain collected in Florida onto the CS background (Adhikari et al., 2021; Kavi et al., 2014). These two strains were raised at 18°C or 29°C for one complete generation (adult to adult) with 12 h: 12 h light:dark photoperiods. Adult flies for each genotype-by-temperature (G×T) combination were kept in 30 cm^3^ cages with containers of 1:1 sugar:nonfat-dry-milk and *ad libitum* water. Eggs were collected in a standard medium consisting of wheat bran, calf manna, wood chips, yeast, and water (Hamm et al., 2009). The same medium was also used for larval development. All stages of the life cycle (larval development, pupation, and adult emergence) were maintained at a constant temperature (either 18°C or 29°C) for one complete generation. Collecting flies after one generation ensured at least one full egg-to-adult cycle at the appropriate temperature.

We sampled 26 unmated lab-raised male flies (within 24 hours of emergence from pupae) from each of the G×T combinations. Newly emerged flies were briefly anesthetized with low pressure CO_2_ using a fly pad and forceps that had been cleaned with 70% ethanol. Males were placed into chilled ethanol on ice and frozen at −80℃. The fly pad and forceps were cleaned with ethanol before and after each G×T sampling to minimize cross-contamination between G×T conditions.

### Wild-caught house flies

Adult flies were collected at multiple time points during 2020 and 2021 from Bastrop, Washington and Johnson Counties in the state of Texas, USA (Table 1). Flies from Bastrop were collected near chickens, goats, and donkeys, whereas flies from Washington were collected within stables containing two horses. House flies were captured with a sweep net, put into ethanol in 1.5 mL tubes, and kept on ice during transport to the laboratory. Wild-caught flies were subsequently stored at −80°C until DNA extraction. Wild-caught flies were sorted into sexes and species (with the goal of identifying male *M. domestica* individuals) based on morphological features (Thomsen, 1938). We only included male flies in our study because we aimed to investigate the effects of Y^M^ and III^M^ on fly-associated microbiomes. We collected a total of 87 wild male flies from the two sites in Texas (Table 1).

**Table 1:** Collection time and sites of wild house flies.

| Collection site | coordinates | Collection date | Number of flies |
| --- | --- | --- | --- |
| Bastrop County | 30.10° N, 97.31° W | August 2020 | 7 |
|  |  | July 2021 | 14 |
| Washington County | 30.21° N, 96.41° W | November 2020 | 15 |
|  |  | May 2021 | 50 |
| Johnson County | 32.37° N, 97.39° W | November 2020 | 1 |

### DNA extraction and 16S rRNA gene amplicon sequencing

DNA was extracted in a dedicated clean work laboratory area using an alkaline lysis method (Supplemental Methods) (Truett et al., 2000). The quantity and purity of DNA was analyzed using a NanoDrop spectrophotometer. Qualified samples were stored at −80°C until being used for Illumina library preparation. The V3-V4 region of DNA was amplified using the QIAseq 16S Region Panels kit (QIAGEN) following the manufacturer’s instructions and 16S libraries were sequenced on an Illumina NextSeq 500 at the University of Houston (UH) Seq-N-Edit core. Sequenced reads were demultiplexed and paired end reads were matched using the standard Illumina pipeline.

### Bacterial 16S rRNA data processing and analysis

We analyzed the 16S sequence data using the QIIME-2 pipeline (Bolyen et al., 2019). Paired-end reads were denoised using the DADA2 plugin to generate amplicon sequence variants (ASVs), resulting in a feature table containing the frequency of each ASV per sample. Truncation parameters were selected based on quality profiles. For lab-reared flies, reads were truncated at 276 bp (forward) and 246 bp (reverse), with a 1 bp trim applied to reverse reads. For wild-caught flies, reads were truncated at 275 bp (forward) and 256 (reverse) with the same trimming applied. For the wild-caught dataset, a total of 88 samples were initially processed for denoising, including one lab fly sample and one sample from Johnson County. These two samples were excluded to ensure dataset consistency resulting in 86 wild-caught fly samples for downstream analysis.

To assess the sufficiency of sequencing depth, alpha rarefaction analysis was performed separately for each data set. For lab-grown flies, rarefaction curves were evaluated up to 7000 sequences per sample whereas for wild flies, curves were assessed up to 7500 sequences per sample. To standardize sequencing depth across samples, lab-reared flies were rarefied to 1,628 sequences per sample, while wild-caught samples were rarefied to 975 sequences per sample. These rarefaction criteria resulted in the exclusion of eight lab-reared samples (retaining 96/104 samples, 92.31%) and six wild-derived samples (retaining 80/86 samples, 93.02%). The SILVA database (Quast et al., 2013) was used to assign sequences to taxonomic levels according to a 97% sequence identity threshold (Stackebrandt & Goebel, 1994). Alpha diversity and beta diversity were calculated for all samples using observed taxa. Statistical comparisons of diversity metrics were performed using ANOVA and PERMANOVA tests. In addition, PERMDISP was used to test which condition has more dispersion or variability within a group. The differential relative abundance of particular taxa was determined by ANCOM (Mandal et al., 2015), which identifies taxa that are significantly different between groups based on the number of pairwise log-ratio comparisons that show significant differences. Additional details are provided in the Supplemental Materials.

## Results

We characterized the bacterial communities associated with 104 laboratory-reared and 86 wild-caught house flies using 16S rRNA amplicon sequencing (Supplemental Table S1). We obtained 816,392 sequencing reads from the 104 lab-reared flies and 757,417 reads from the 86 wild-caught flies, which were assigned to 3,909 and 4,050 amplicon sequence variants (ASVs), respectively, using QIIME 2 (Supplemental Table S2). However, after rarefaction, we retained 96 lab-reared samples and 80 wild-caught samples for downstream analysis. Rarefaction curves demonstrate that we had sufficient sequencing depth to measure the taxonomic diversity in our samples (Supplemental Figures S1 and S2).

### Temperature affects the house fly microbiome

We used multiple alpha diversity metrics to quantify bacterial community diversity within each lab-reared house fly microbiome (Figure 1; Supplemental Table S3). Most of the alpha diversity metrics did not significantly differ across temperatures or genotypes (Supplemental Table S3). However, Faith’s phylogenetic diversity metric was significantly affected by temperature (*p*<0.0001), with higher alpha diversity at 29°C than 18°C (Figure 1). These results are not affected by our sequencing depth (Supplemental Figure S1). In addition, Pielou’s evenness was significantly affected by a G×T interaction (p = 0.043), with higher diversity in III^M^ flies at 29°C.

**Figure 1.**
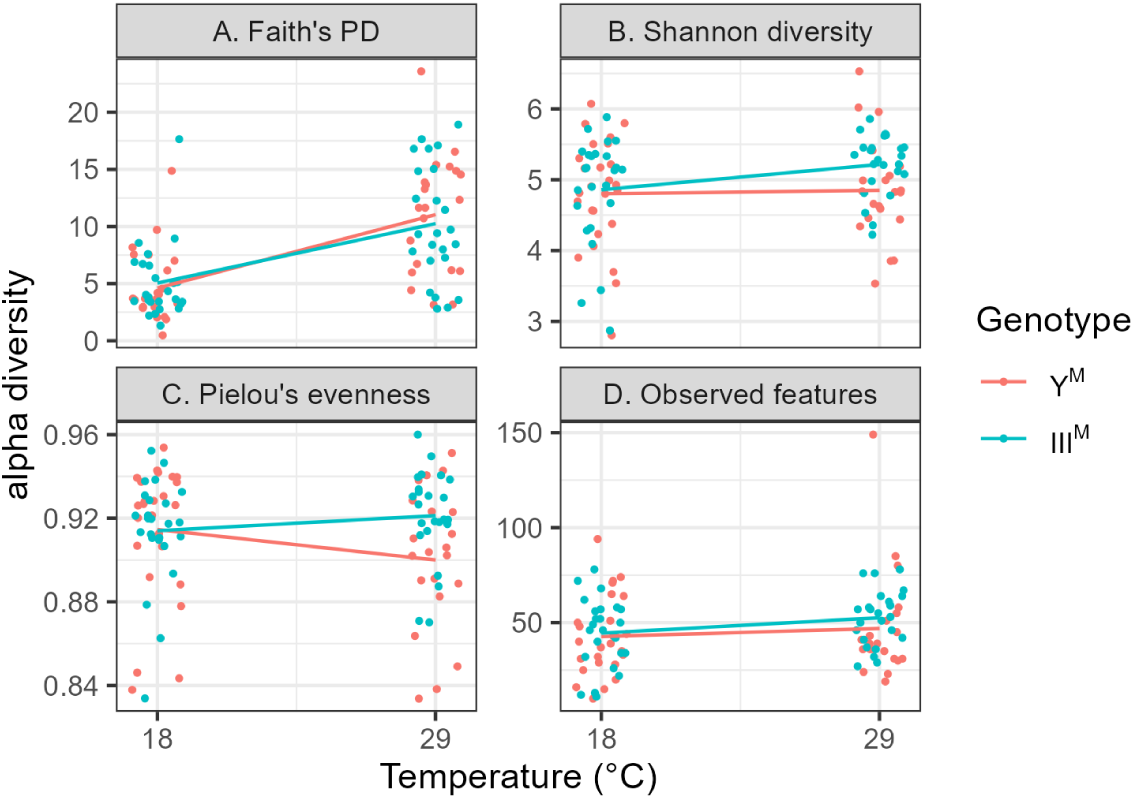
Alpha diversity metrics of lab-reared house fly microbiome. Each dot is an individual fly with either the III^M^ (salmon) or Y^M^ (cyan) genotype raised at 18°C or 29°C. Lines connect the means of each genotype at the two temperatures.

We next used multiple beta diversity metrics to compare microbial community composition across lab-reared flies (Figure 2). PERMANOVA analyses revealed a significant effect of temperature on microbiome composition for all four beta diversity metrics that we applied (Supplemental Table S4). In contrast, genotype affected inter-fly microbiome differences only when using the weighted UniFrac and Bray-Curtis metrics. We also found significant G×T interactions on between-fly diversity using the Jaccard (p=0.035) and weighted UniFrac (p=0.003) metrics, and the effects were close to significant by unweighted UniFrac (p=0.067) and Bray-Curtis (p=0.068). In addition, PCoA revealed that the phylogenetic distance-based metrics explained more bacteriome variation among flies than the non-phylogenetic metrics (Figure 2). Specifically, weighted UniFrac and unweighted UniFrac showed higher percentages of variation explained by the first two Principal Coordinates (PCoA), indicating strong separation of bacterial communities based on phylogenetic relationships. Furthermore, PERMDISP analysis revealed more dispersion (i.e., more between-fly heterogeneity) in flies raised at 29°C than flies at 18°C (Supplemental Figure S3). These effects were significant by weighted UniFrac (p=0.008), Jaccard (p=0.04), and Bray-Curtis (p=0.05) distance metrics (Supplemental Table S5). In contrast, genotype did not affect the variability significantly (Supplemental Table S5; Supplemental Figure S4). Taken together, our results suggest that at 29°C, not only do individual house flies harbor more phylogenetically diverse microbiomes, but the composition of those microbiomes was more variable between flies.

**Figure 2.**
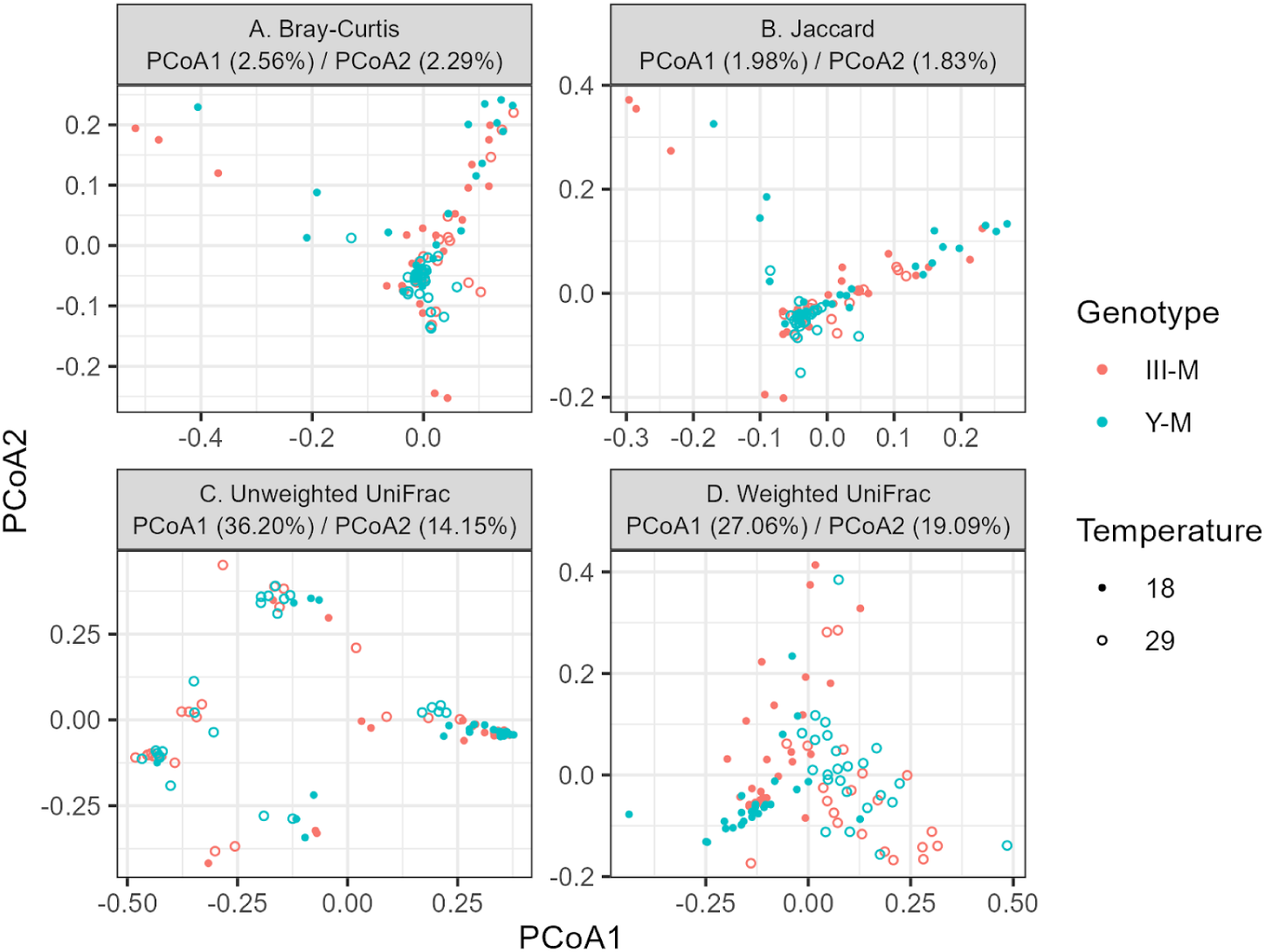
Beta diversity metrics of lab-reared house fly microbiome. The X and Y axes report the principal coordinate axes (PCoA). Each point represents an individual fly with either a III^M^ (salmon) or Y^M^ (cyan) genotype raised at either 18°C (solid) or 29°C (hollow).

We then identified the bacterial taxa responsible for the differences in communities between III^M^ and Y^M^ house flies reared at 29°C and 18°C. Proteobacteria was the most abundant phylum across both genotypes and temperature conditions, and Bacteroidota and Firmicutes were also consistently present in varying proportions (Supplemental Figure S4). However, the overall phylum composition was mostly similar across temperatures and genotypes. Consistent with this observation, ANCOM analysis did not detect any differentially abundant phyla between genotypes or temperatures. In contrast, we observed inter-genotype and inter-temperature differences at the class, order, family, and genus levels (Supplemental Tables S6-S9). Higher abundance of Gammaproteobacteria (a class within Proteobacteria) was observed at 18°C, while Clostridia and Bacilli (classes within Firmicutes) were more abundant at 29°C. The order Haloplasmatales and class Haloplasmataceae were more abundant in Y^M^ males than III^M^ individuals at 18°C (Supplemental Table S6), whereas Xanthomonadales and Xanthomonadaceae were more abundant in Y^M^ than III^M^ at 29°C (Supplemental Table S7).

We observed many differences at the genus level. For example, *Chryseobacterium*, *Stenotrophomonas*, and *Dietzia* were more abundant in Y^M^ males than III^M^ males at both temperatures (Supplemental Tables S6 and S7). There were also several bacterial genera that significantly varied in relative abundance between temperature treatments regardless of genotype (Supplemental Tables S8 and S9). For example, *Paenibacillaceae*, *Brachybacterium*, and *Proteus* were more abundant in flies reared at 18°C, but most differentially abundant genera were genotype-specific. Within Y^M^ flies, the majority of differentially abundant taxa exhibited higher abundance at 18°C than at 29°C, including *Leucobacter, Myroides, Haloplasma,* and *Advenella.* However, other taxa—such as *Chryseobacterium, Enterococcus, Ochrobactrum, Pseudomonas,* and *Stenotrophomonas*—were more prevalent at 29°C. In the III^M^ genotype, a similar trend was observed, where most different abundant taxa were more prevalent at 18°C, including *Amphibacillus,* uncultured *Bacillaceae, Pseudochrobactrum, Bordetella,* and *Morganella*. In contrast, a few genera, such as *Glutamicibacter*, *Zymobacter*, and unclassified *Erwiniaceae*, were more abundant at 29°C.

### Differences in microbial communities associated with wild house flies at two collection sites

We sampled wild male house flies from two counties (Bastrop and Washington) in Texas, USA, in 2020 and 2021. Our PCR genotyping assay identified that all flies carried III^M^, and no flies carried Y^M^. We therefore could not compare between genotypes in our wild-caught flies. We analyzed data from both collection years together because we had smaller sample sizes in 2020 and collected in different months at the two sites (Table 1), which limited our ability to analyze both site and collection date in a single statistical model. Phylogenetic diversity, Shannon diversity, and observed features were significantly different between flies from the Bastrop and Washington collection sites (Figure 4; Supplemental Table S10). In all cases, flies collected from Bastrop County had higher alpha diversity than those collected in Washington County (Figure 4). We also detected significant differences in beta diversity between flies collected from the two counties using both the unweighted and weighted UniFrac distances, but not the Bray-Curtis and Jaccard metrics (Figure 5; Supplemental Table S11). However, PERMDISP analysis did not show any significant location effects on the dispersion or variability within each location (Supplemental Table S12).

Taxonomic analysis of the wild caught flies revealed additional differences in the microbiomes across sites. Proteobacteria was the most dominant phylum in the wild caught flies, consistently representing a major proportion of the microbial community across all samples (Supplemental Figure S6). Bacteroidota and Firmicutes were also prominent in several samples, although their relative abundance varied. In 2021, there were more differentially abundant taxa at each taxonomic level between Bastrop and Washington because of higher sample size (i.e., increased power) compared to 2020 (Table 1; Supplemental Tables S13 and S14). For example, Fusobacteriota and Desulfobacterota were significantly enriched in Bastrop relative to Washington County in 2021 (Supplemental Table S14) but not in 2020 (Supplemental Table S13). At the family level, Bacteroidaceae, Ruminococcaceae, Rikenellaceae, Lactobacillaceae, Fusobacteriaceae, Tannerellaceae, and Peptostreptococcaceae showed higher relative abundance in Bastrop samples. Similarly, at the genus level, *Bacteroides, Lactobacillus, Romboutsia*, Rikenellaceae_RC9_gut_group, *Escherichia-Shigella, Megamonas, Alistipes, Alloprevotella*, and an unidentified genus were also significantly more abundant in Bastrop (Figure 6). These results strongly indicate distinct microbial communities composition associated with flies from each collection site.

## Discussion

We evaluated how genotype and environment affect the bacterial community composition of lab-raised and wild-caught male house flies. Temperature had a much greater effect on the lab-raised house fly microbiome than genotype (Figures 1 and 2). Specifically, at a warmer temperature (29°C), individual male house flies harbored more phylogenetically diverse microbiomes, and the composition of those microbiomes was more variable between flies than flies raised at a cooler temperature (18°C). Consistent with large environmental effects, house flies collected from two locations <120 km apart differed in their average within fly bacterial diversity and their between fly bacterial phylogenetic diversity (Figure 4 and 5), despite all possessing the same III^M^ genotype.

### Temperature affects the house fly microbiome

Our findings revealed that temperature exerts an impact on the house fly microbiome, with effects that are most evident when considering phylogenetic relationships among bacterial taxa. For example, Faith’s phylogenetic diversity was greater in flies raised at 29°C than those raised at 18°C, but no non-phylogenetic alpha diversity metrics differed between temperatures (Figure 1). This pattern is opposite from what has been previously characterized in *D. melanogaster*, where increasing temperature decreases alpha diversity (Moghadam et al., 2018). In addition, phylogenetic beta diversity metrics (weighted and unweighted UniFrac) explained more variation among flies than non-phylogenetic metrics (Figure 2), but temperature affected beta diversity regardless of the metric. For most beta diversity metrics, the between fly microbiome diversity was greater at 29°C than 18°C. Our results therefore demonstrate that increasing temperature causes the house fly microbiome to become more diverse within each individual fly and more variable across flies.

Temperature affects the relative abundance of several bacterial genera in the house fly microbiome in ways that are consistent with what has been previously observed in those genera. For example, *Enterococcus* and *Stenotrophomonas* were more abundant in house flies raised 29°C, consistent with those genera thriving at warmer temperatures (Panagea & Chadwick, 1996; Wang et al., 2024). A similar enrichment of heat-tolerant bacterial taxa was documented in mosquitoes upon exposure to elevated temperature (Onyango et al., 2020). Previous studies also found that the microbiomes of insects and other ectotherms reared at high temperatures experienced an increase in the abundance of Proteobacteria (Horváthová et al., 2019; Y.-F. Li et al., 2018; Moghadam et al., 2018). Of the eight genera enriched at 29°C in either house fly genotype, five were Proteobacteria (*Ochrobactrum*, *Pseudomonas*, *Stenotrophomonas*, *Zymobacter*, and *Erwiniaceae*). In contrast, 5/14 genera enriched at 18°C were Proteobacteria, whereas six were in the phylum Firmicutes. This modest, but not significant, over-representation of Proteobacteria amongst the genera enriched at 29°C is consistent with previous observations in other ectotherms (Horváthová et al., 2019; Y.-F. Li et al., 2018; Moghadam et al., 2018).

There was also a G×T effect on both alpha and beta diversity using some of the metrics, providing evidence for a genotype effect on the microbiome that depends on temperature. For example, Pielou’s evenness was reduced in III^M^ males at 29°C compared to Y^M^ (Figure 1). We also found that most temperature-dependent changes in the abundance of genera were limited to one house fly genotype (Figure 3). This provides additional evidence for the host genotype mediating how the house fly microbiome responds to temperature. The III^M^ chromosome is found in southern populations and confers greater heat tolerance and preference for warmer temperatures, while Y^M^ is associated with cold-adapted traits (Delclos et al., 2021). It remains to be determined, however, if the differences in microbiomes at the warmer temperature are causally associated with any behavioral or physiological differences between house fly genotypes.

**Figure 3.**
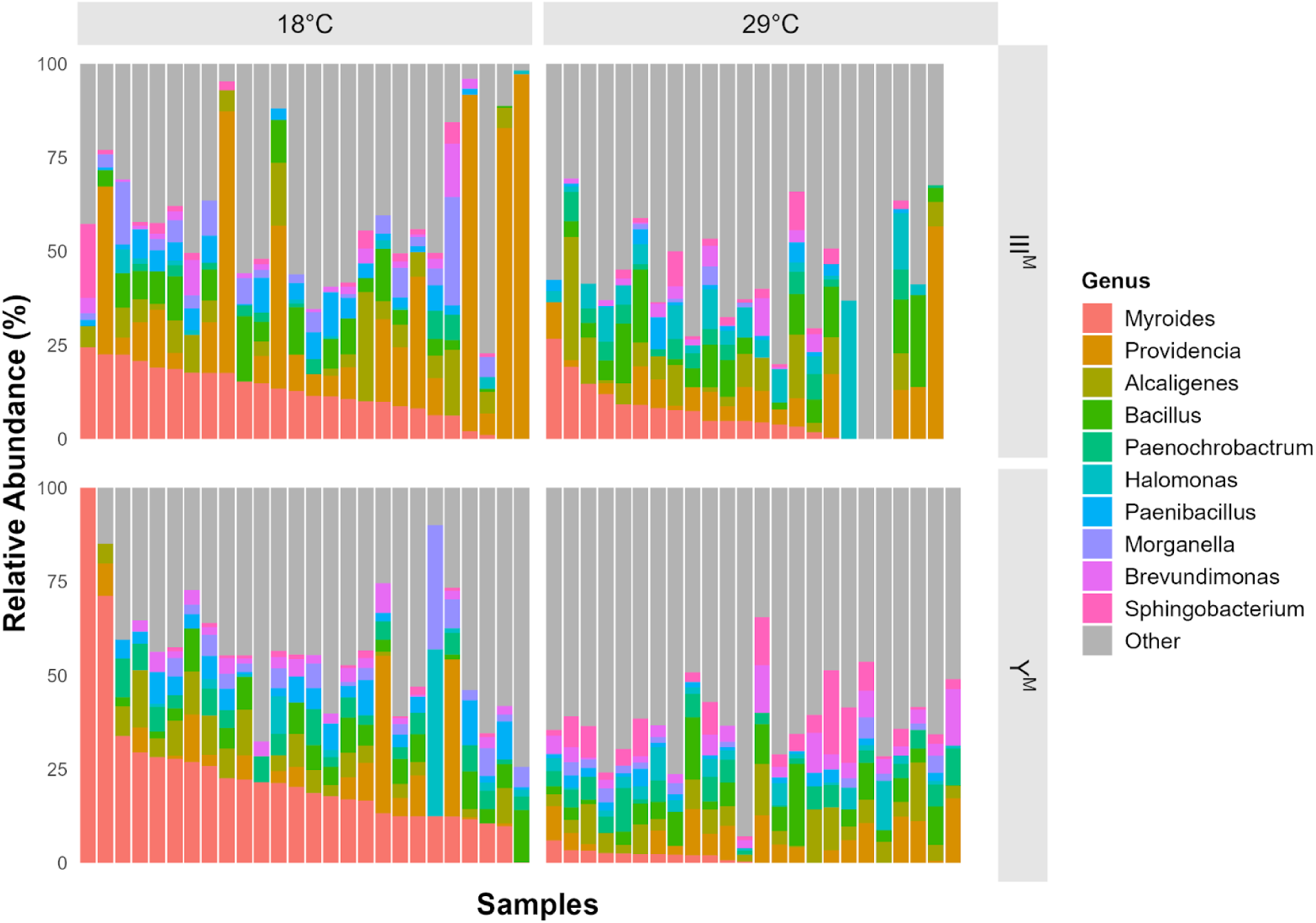
Relative abundance of the top 10 bacterial genera across samples across genotype and temperature. Stacked bar plots show the relative abundance (%) of the 10 most abundant bacterial genera across individual samples of *Musca domestica* lab flies. Genera outside the top 10, as well as unclassified/uncultured taxa, are grouped as “Other” (gray). Samples are faceted by genotype (III^M^ and Y^M^) and rearing temperature (18°C and 29°C). Each bar represents a single sample, and the color legend indicates genus identity.

We further identified specific bacterial taxa that differed in abundance between genotypes in ways that depended on temperature. For example, at 18°C, we observed multiple taxa within Firmicutes that were more abundant in Y^M^ males than III^M^, including Clostridia and Bacilli (Haloplasmatales: Haloplasmataceae). At 29°C, Proteobacteria were more abundant in Y^M^ males, particularly Gammaproteobacteria, Xanthomonadales, and Xanthomonadacea. In contrast, specific genera (e.g., *Chryseobacterium*, *Dietzia*, and *Stenotrophomonas*) were more abundant in Y^M^ males regardless of temperature. Our results therefore suggest that, while some genera are consistently enriched in one genotype, higher-level taxonomic structure differentially responds to genotype in a way that depends on temperature.

### Microbiome variation across wild house flies and differences from lab-reared flies

We sampled wild house flies from two locations that differed in the large animals present. Specifically, the Bastrop collection site had a more diverse collection of livestock and poultry (chickens, goats, and donkeys) than the Washington site (a horse stable). While previous work has identified differences in house fly microbiomes across locations (Bahrndorff et al., 2020; Neupane et al., 2023; Neupane & Nayduch, 2022), other studies have shown that most of the variation in house fly microbiome richness and diversity is observed between individuals, not locations (Bahrndorff et al., 2017). Consistent with the latter observation, the microbiomes associated with house flies at both of our collection sites were largely overlapping (Figure 5 and 6). Nonetheless, we did identify important differences between sites. For example, the microbiomes of flies from the Bastrop site had greater alpha diversity than those collected in Washington County (Figure 4), suggesting that an elevated diversity of domesticated animals leads to a more diverse house fly microbiome. Other work has shown that house flies have a stable internal bacterial microbiota across populations, but the external bacterial microbiota vary across geographical location and habitat (Park et al., 2019). We sampled the house fly microbiome without distinguishing between external and internal bacteria (although we did wash the exterior with ethanol), and we therefore lack the resolution to test for differences between surface and gut communities. Nevertheless, our comparisons across sampling sites provides additional evidence that the environment affects the house fly microbiome more than genotype.

**Figure 4.**
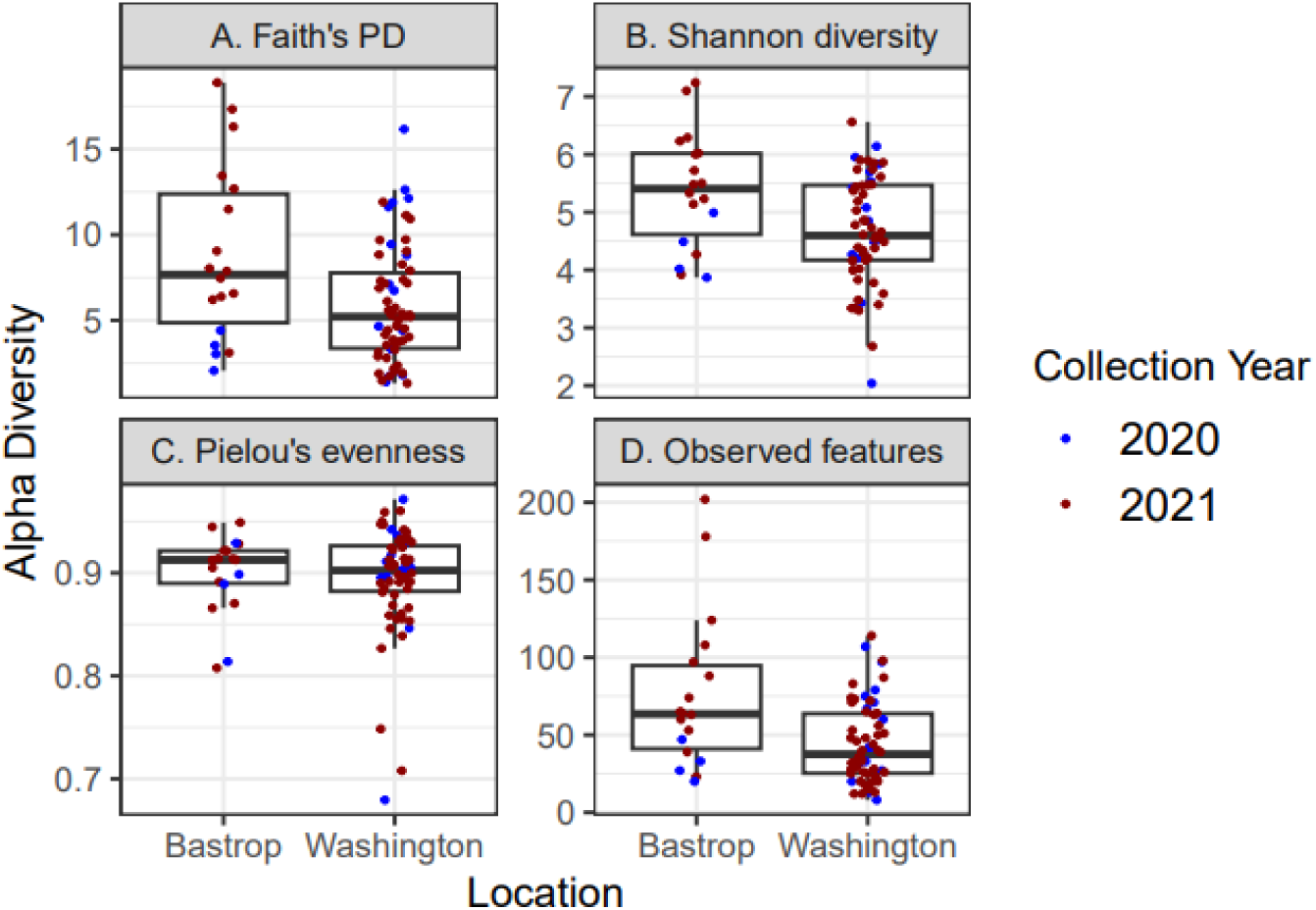
Alpha diversity metrics of wild-caught house fly microbiome. Each dot represents the microbial community of an individual fly collected from Bastrop or Washington County in either 2020 (blue) or 2021 (red). Boxplot shows the median and quartile in each county at each collection date.

**Figure 5.**
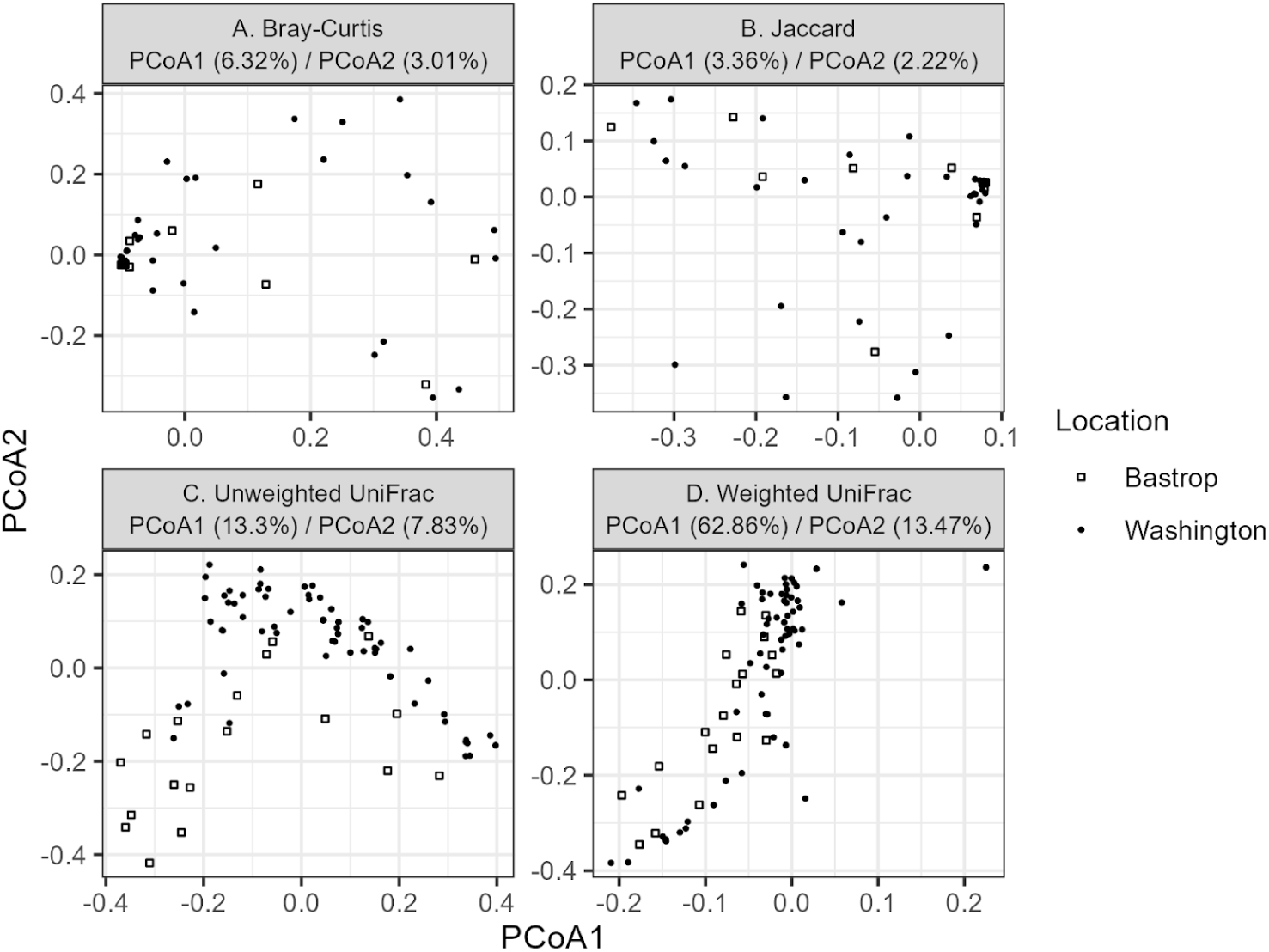
Beta diversity metrics of wild-caught fly microbiome. The X and Y axes report the principal coordinate axes (PCoA) for four separate beta diversity metrics. Each point represents an individual fly collected in either Bastrop (empty square) or Washington (solid circle).

**Figure 6.**
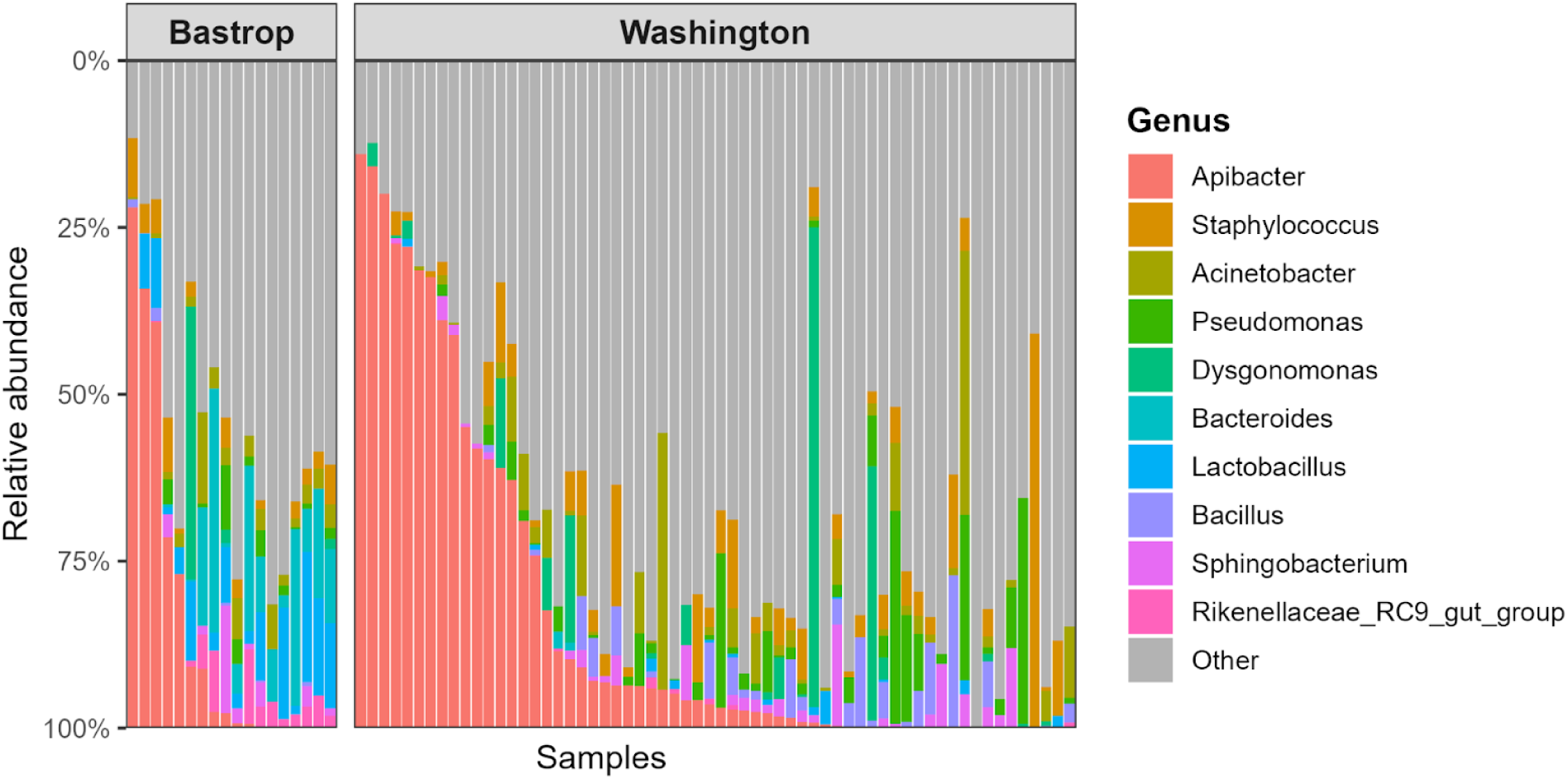
Relative abundance of the top 10 bacterial genera across house fly samples. Stacked bar plots showing the relative abundance of the 10 most abundant bacterial genera across samples from different collection locations. Genera outside the top 10, as well as unclassified/uncultured taxa, were grouped as “Other.” Each bar represents one sample, with the y-axis scaled to relative abundance (proportion of total reads per sample). Colors indicate bacterial genera, ordered by total abundance across all samples.

We identified specific bacterial taxa that differed in abundance between Bastrop and Washington in a way that was consistent with the livestock and poultry present at each site. For example, Fusobacteriota were enriched in the Bastrop samples, and this phylum is associated with animal mucosal habitats, including the oral and gastrointestinal systems (Nagaraja et al., 2005; Pignatelli et al., 2023; Schwarz et al., 2023). In addition, many of the taxa enriched in Bastrop flies were members of the families Bacteroidaceae and Ruminococcaceae, well-known constituents of the gastrointestinal microbiota of livestock and poultry (Burrows et al., 2025; Y. Li et al., 2022; Mahayri et al., 2022; Yue et al., 2024; R. Zhang et al., 2020; Zhuang et al., 2020). Similarly, Desulfobacterota was more abundant in Bastrop flies compared to those from Washington, consistent with previous findings that Desulfobacterota are enriched in environments impacted by agricultural animal husbandry (Muyyarikkandy et al., 2023; X. Zhang et al., 2022). Prior work has shown that house flies share almost all bacterial taxa with the manure present at their collection sites (Neupane, Park, et al., 2024; Neupane, Talley, et al., 2024). The higher diversity of livestock and poultry at the Bastrop site appear to have been important factors shaping the house fly microbiome, and the differential abundance of taxa across sites may provide insight into how local agricultural practices can influence the microbial communities of insects.

We also observed differences between the wild house flies and lab-reared flies that are consistent with previous work. In particular, wild house flies are expected to have more diverse microbiomes than lab reared flies, but some taxa are expected to be shared regardless of a fly’s origin (Voulgari-Kokota et al., 2022). Consistent with the first expectation, wild-caught house flies in our samples exhibited a slightly higher number of distinct taxonomic units than lab-grown flies, despite fewer wild-caught samples (Supplemental Table S2). The latter expectation was also supported—Proteobacteria was the most dominant phylum in both the lab-reared and wild-caught house flies, representing a major proportion of the microbial community across all samples. However, we observed differences in the most abundant bacterial genera between lab-reared and wild-caught house flies. In the lab-reared flies, *Providencia* and *Myroides* were the most common genera (Figure 3), in accordance with prior observations of laboratory house flies (Voulgari-Kokota et al., 2022). In wild-caught flies, *Apibacter*, *Staphylococcus*, and *Acinetobacter* were the most abundant genera, but their dominance varied substantially across samples (Figure 6). *Apibacter* is found in bee guts, and it is thought to be a beneficial symbiont in bumble bees (Kwong & Moran, 2016; Mockler et al., 2018; Praet et al., 2016). Species in the genera *Staphylococcus* and *Acinetobacter*, in contrast, are important human pathogens (Peleg et al., 2008; Tong et al., 2015), providing additional evidence that wild house flies could to serve as vectors for enteropathogens (Nayduch et al., 2023).

### Dysbiosis and thermal stress

Lastly, we aimed to test if shifting a warm adapted animal to a cooler temperature disrupted the microbiome. We did indeed observe that temperature substantially affected the house fly microbiome, but these temperature differences are not necessarily evidence of a disrupted microbiome. Dysbiosis is a conventional term for microbiome disruption (Levy et al., 2017). Specifically, dysbiotic microbiomes are expected to be different from each other, while healthy microbiomes are more uniform across samples (Zaneveld et al., 2017). Dysbiosis is thus predicted to result in greater beta diversity, which is thought to emerge via the dominance of atypical bacterial taxa unique to each sample. The dominance of these atypical taxa will result in reduced alpha diversity within dysbiotic individuals. While we did observe reduced alpha diversity at a lower temperature in house fly, we also saw greater beta diversity at the higher temperature. We therefore conclude that low temperature does not cause dysbiosis in a warm-adapted host. Instead, higher temperatures increase both alpha and beta diversity in house flies, suggesting that we did not expose the flies to heat stress at 29°C.

## Acknowledgements

This work was supported by the National Science Foundation grant DEB 1845686 (to RPM), the Texas EcoLab (to KA), and Sigma Xi Grant-in-Aid of Research G2020100194691045 (to KA). Jyoti Lama assisted with collecting the wild house flies, and Sara Loetzerich assisted with rearing the laboratory flies.

## Data Availability

Sequence data are available from the NCBI SRA at BioProjects PRJNA1415840 and PRJNA1415960. Analysis methods are available at https://doi.org/10.18738/T8/Y2I9HK.

## Supplemental Methods

DNA was extracted from single flies using an alkaline lysis method. Each fly was homogenized in 100 uL of solution A (100mM TrisHCl, 20mM EDTA, 100mM NaCl, and 0.5% SDS at of at pH 7.5 in water) and incubated at 95°C for 30 minutes. 160 uL of solution B (1.43 M potassium acetate and 4.29 M LiCl in water) was added to the tube, the solution in the tube was mixed, and the tube was placed in ice for 10 minutes. The tube was then vortexed and centrifuged at maximum speed for 15 minutes. The supernatant was transferred to a new tube, and 120 uL of isopropanol was added. The sample was mixed thoroughly and centrifuged again for 15 minutes. The supernatant was discarded, the remaining pellet was washed with 70% ethanol, and the sample was centrifuged for 3 minutes. The supernatant was discarded, and the remaining pellet was allowed to air dry for 5-10 minutes. The pellet was then resuspended in 50ul of nuclease free water.

## Supplemental Tables

**Supplemental Table S1:** Summary of demultiplexed 16S rRNA sequencing reads from lab-grown and wild-caught flies (these include 87 wild-caught flies and 1 lab flies). Reported values include minimum, maximum, mean, and median read pairs per sample, along with the total number of reads recovered.

| House fly strains |  | Read pairs |
| --- | --- | --- |
| Lab-grown flies | Minimum | 8786 |
|  | Median | 83346.5 |
|  | Mean | 122082.490385 |
|  | Maximum | 2406115 |
|  | Total | 12696579 |
| Wild-caught flies | Minimum | 5350 |
|  | Median | 72863 |
|  | Mean | 78996.659091 |
|  | Maximum | 217209 |
|  | Total | 6951706 |

**Supplemental Table S2:** Summary of feature table statistics for lab-grown and wild-caught flies. Metrics include the number of samples, total number of features (ASVs), and sequencing frequencies across all samples.

| House fly strains | Metric | Sample |
| --- | --- | --- |
| Lab-grown flies | No. of samples | 104 |
|  | No. of features | 3909 |
|  | No. of frequency | 816,392 |
| Wild-caught flies | No. of samples | 86 |
|  | No. of features | 4,050 |
|  | No. of frequency | 757,417 |

**Supplemental Table S3.** ANOVA test to examine the effects of G×T on alpha diversity of lab-grown fly microbiome.

| Alpha diversity metric |  | PR(>F) |
| --- | --- | --- |
| Faith's phylogenetic diversity | Temperature (T) | **<0.001\*** |
|  | Genotype (G) | 0.878922 |
|  | G*T | 0.47321 |
| Pielou's evenness | Temperature (T) | 0.568782 |
|  | Genotype (G) | 0.117289 |
|  | G*T | **0.04372\*** |
| Shannon diversity | Temperature (T) | 0.144091 |
|  | Genotype (G) | 0.156104 |
|  | G*T | 0.278245 |
| Observed features | Temperature (T) | 0.146393 |
|  | Genotype (G) | 0.399524 |
|  | G*T | 0.656914 |

**Supplemental Table S4.** Adonis test (PERMANOVA) to examine the effects of G ×T on beta diversity of lab-grown fly microbiome.

| Beta diversity metric |  | PR(>F) |
| --- | --- | --- |
| Unweighted UniFrac distance | Temperature (T) | **0.001\*** |
|  | Genotype (G) | 0.155 |
|  | G×T | 0.067 |
| Bray-Curtis distance | Temperature (T) | **0.001\*** |
|  | Genotype (G) | **0.014\*** |
|  | G×T | 0.068 |
| Jaccard distance | Temperature (T) | **0.001\*** |
|  | Genotype (G) | 0.160 |
|  | G×T | **0.035\*** |
| Weighted UniFrac distance | Temperature (T) | **0.002\*** |
|  | Genotype (G) | **0.001\*** |
|  | G×T | **0.003\*** |

**Supplemental Table S5.** PERMDISP to examine the effects of G ×T on dispersion of lab-grown fly microbiome.

| Beta diversity metric |  | PR(>F) |
| --- | --- | --- |
| Unweighted UniFrac distance | Temperature (T) | 0.66 |
|  | Genotype (G) | 0.15 |
|  | G×T | 0.58 |
| Bray-Curtis distance | Temperature (T) | **0.05** |
|  | Genotype (G) | 0.07 |
|  | G×T | 0.43 |
| Jaccard distance | Temperature (T) | **0.04** |
|  | Genotype (G) | 0.28 |
|  | G×T | 0.65 |
| Weighted UniFrac distance | Temperature (T) | **0.008** |
|  | Genotype (G) | 0.38 |
|  | G×T | **0.012** |

**Supplemental Table S6:**
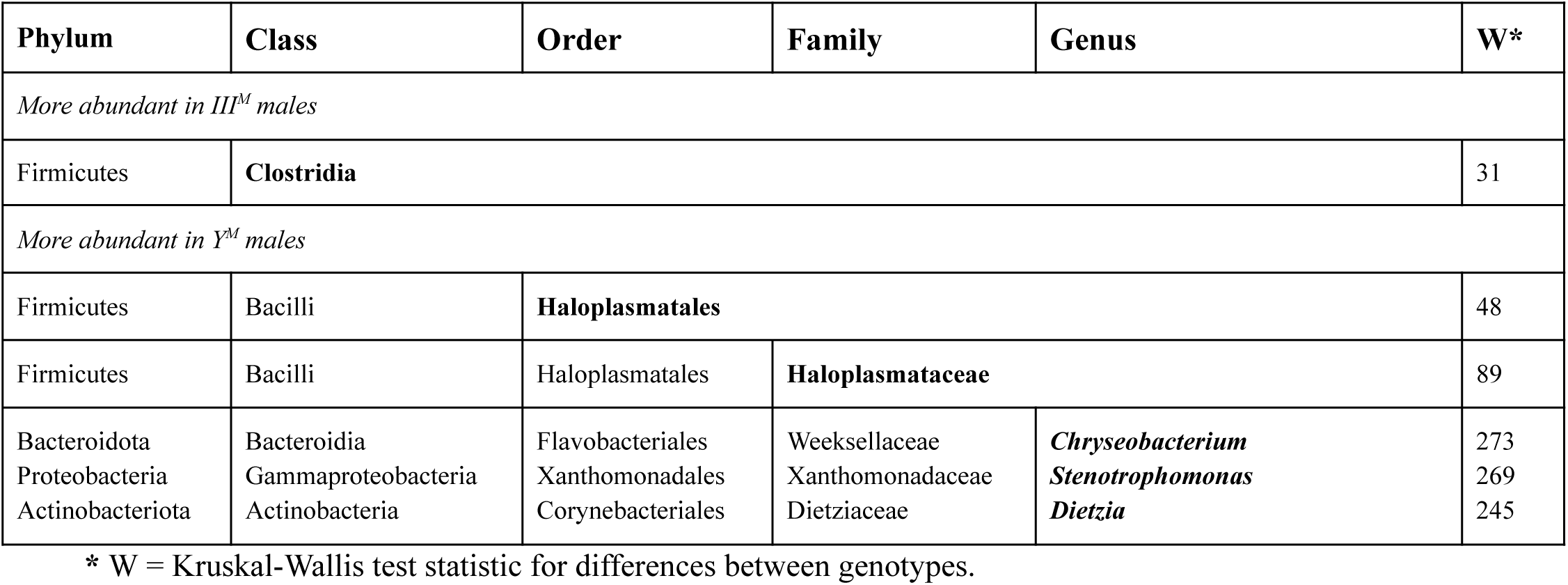
Differentially abundant taxa between the two genotypes at 18°C.

**Supplemental Table S7:** Differentially abundant taxa between the two genotypes at 29°C.

| Phylum | Class | Order | Family | Genus | W* |
| --- | --- | --- | --- | --- | --- |
| *More abundant in Y<sup>M</sup> males* |  |  |  |  |  |
| Firmicutes | **Clostridia** |  |  |  | 31 |
| Proteobacteria | Gammaproteobacteria | **Xanthomonadales** |  |  | 82 |
| Proteobacteria | Gammaproteobacteria | Xanthomonadales | **Xanthomonadaceae** |  | 155 |
| Bacteroidota | Bacteroidia | Flavobacteriales | Weeksellaceae | *Chryseobacterium* | 273 |
| Proteobacteria | Gammaproteobacteria | Xanthomonadales | Xanthomonadaceae | *Stenotrophomonas* | 269 |
| Actinobacteriota | Actinobacteria | Corynebacteriales | Dietziaceae | *Dietzia* | 245 |

\* W = Kruskal-Wallis test statistic for differences between genotypes.

**Supplemental Table S8:** Differentially abundant taxa between two temperatures in Y^M^ males.

| Phylum | Class | Order | Family | Genus | W* |
| --- | --- | --- | --- | --- | --- |
| *More abundant at 18°C* |  |  |  |  |  |
| Firmicutes | Bacilli | **Haloplasmales** |  |  | 77 |
| Bacteroidota | Bacteroidia | **Flavobacteriales** |  |  | 72 |
| Firmicutes | Bacilli | **Paenibacillales** |  |  | 71 |
| Bacteroidota | Bacteroidia | Flavobacteriales | **Flavobacteriaceae** |  | 135 |
| Firmicutes | Bacilli | Haloplasmales | **Haloplasmataceae** |  | 132 |
| Actinobacteriota | Actinobacteria | Micrococcales | **Dermabacteraceae** |  | 131 |
| Actinobacteriota | Actinobacteria | Micrococcales | **Microbacteriaceae** |  | 131 |
| Firmicutes | Bacilli | Paenibacillales | Paenibacillaceae | *Paenibacillaceae* | 231 |
| Bacteroidota | Bacteroidia | Flavobacteriales | Flavobacteriaceae | *Myroides* | 229 |
| Firmicutes | Bacilli | Haloplasmales | Haloplasmataceae | *Haloplasma* | 228 |
| Firmicutes | Bacilli | Bacillales | Bacillaceae | **Uncultured** | 224 |
| Proteobacteria | Gammaproteobacteria | Burkholderiales | Alcaligenaceae | *Advenella* | 220 |
| Actinobacteriota | Actinobacteria | Micrococcales | Microbacteriaceae | *Leucobacter* | 220 |
| Actinobacteriota | Actinobacteria | Micrococcales | Dermabacteraceae | *Brachybacterium* | 218 |
| Proteobacteria | Gammaproteobacteria | Enterobacterales | Morganellaceae | *Proteus* | 214 |
| *More abundant at 29°C* |  |  |  |  |  |
| Firmicutes | **Clostridia** |  |  |  | 30 |
| Proteobacteria | Gammaproteobacteria | **Xanthomonadales** |  |  | 70 |
| Proteobacteria | Gammaproteobacteria | **Pseudomonadales** |  |  | 67 |
| Firmicutes | Clostridia | **Lachnospirales** |  |  | 66 |
| Firmicutes | Bacilli | Lactobacillales | **Enterococcaceae** |  | 125 |
| Proteobacteria | Gammaproteobacteria | Xanthomonadales | **Xanthomonadaceae** |  | 124 |
| Proteobacteria | Gammaproteobacteria | Pseudomonadales | **Pseudomonadaceae** |  | 124 |
| Bacteroidota | Bacilli | Flavobacteriales | **Weeksellaceae** |  | 124 |
| Bacteroidota | Bacteroidia | Flavobacteriales | Weeksellaceae | *Chryseobacterium* | 223 |
| Firmicutes | Bacilli | Lactobacillales | Enterococcaceae | *Enterococcus* | 220 |
| Proteobacteria | Gammaproteobacteria | Pseudomonadales | Pseudomonadaceae | *Pseudomonas* | 219 |
| Proteobacteria | Alphaproteobacteria | Rhizobiales | Rhizobiaceae | *Ochrobactrum* | 218 |
| Proteobacteria | Gammaproteobacteria | Xanthomonadales | Xanthomonadaceae | *Stenotrophomonas* | 213 |

**Supplemental Table S9:** Differentially abundant taxa between two temperatures in III^M^ males.

| Phylum | Class | Order | Family | Genus | W* |
| --- | --- | --- | --- | --- | --- |
| *More abundant at 18°C* |  |  |  |  |  |
| Firmicutes<br>Proteobacteria | Bacilli<br>Gammaproteobacteria | **Paenibacillales**<br>**Enterobacterales** |  |  | 65<br>61 |
| Proteobacteria<br>Firmicutes<br>Actinobacteriota<br>Actinobacteriota | Gammaproteobacteria<br>Bacilli<br>Actinobacteria<br>Actinobacteria | Enterobacterales<br>Paenibacillales<br>Micrococcales<br>Micrococcales | **Morganellaceae**<br>**Paenibacillaceae**<br>**Dermabacteriaceae**<br>**Microbacteriaceae** |  | 116<br>115<br>115<br>113 |
| Proteobacteria<br>Firmicutes<br>Firmicutes<br>Proteobacteria<br>Proteobacteria<br>Firmicutes<br>Actinobacteriota<br>Proteobacteria<br>Proteobacteria | Alphaproteobacteria<br>Bacilli<br>Bacilli<br>Gammaproteobacteria<br>Gammaproteobacteria<br>Bacilli<br>Actinobacteria<br>Gammaproteobacteria<br>Gammaproteobacteria | Rhizobiales<br>Paenibacillales<br>Bacillales<br>Burkholderiales<br>Enterobacterales<br>Bacillales<br>Micrococcales<br>Enterobacterales<br>Enterobacterales | Rhizobiaceae<br>Paenibacillaceae<br>Bacillaceae<br>Alcaligenaceae<br>Morganellaceae<br>Bacillaceae<br>Dermabacteraceae<br>Morganellaceae<br>Enterobacteriaceae | *Pseudochrobactrum*<br>*Paenibacillaceae*<br>**Uncultured**<br>*Bordetella*<br>*Morganella*<br>*Amphibacillus*<br>*Brachybacterium*<br>*Proteus*<br>- | 203<br>203<br>202<br>202<br>201<br>200<br>194<br>192<br>191 |
| *More abundant at 29°C* |  |  |  |  |  |
| Proteobacteria<br>Proteobacteria<br>Firmicutes | Gammaproteobacteria<br>Gammaproteobacteria<br>Bacilli | **Oceanospirillales**<br>**Pseudomonadales**<br>**Staphylococcales** |  |  | 64<br>60<br>55 |
| Proteobacteria<br>Proteobacteria | Gammaproteobacteria<br>Gammaproteobacteria | Oceanospirillales<br>Enterobacterales | **Halomonadaceae**<br>**Erwiniaceae** |  | 107<br>107 |
| Proteobacteria<br>Proteobacteria<br>Actinobacteriota | Gammaproteobacteria<br>Gammaproteobacteria<br>Actinobacteria | Oceanospirillales<br>Enterobacterales<br>Micrococcales | Halomonadaceae<br>Erwiniaceae<br>Micrococcaceae | *Zymobacter*<br>-<br>*Glutamicibacter* | 192<br>184<br>177 |

\* W = Kruskal-Wallis test statistic for differences between temperatures

**Supplemental Table S10.**
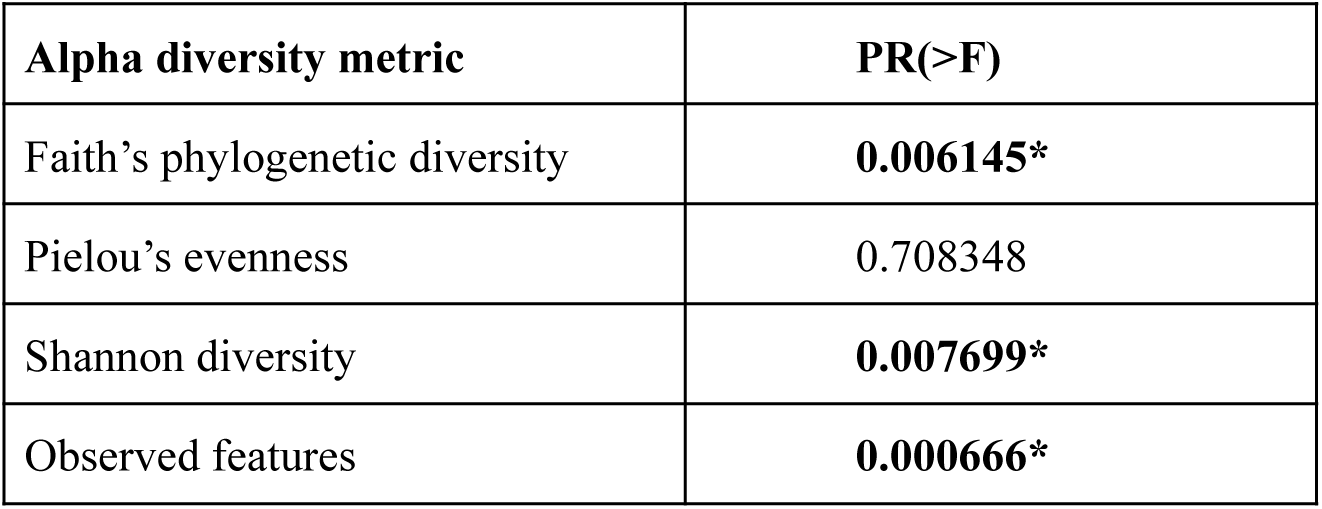
ANOVA test to examine the effects of location on alpha diversity of wild fly microbiome.

| Alpha diversity metric | PR(>F) |
| --- | --- |
| Faith's phylogenetic diversity | **0.006145\*** |
| Pielou's evenness | 0.708348 |
| Shannon diversity | **0.007699\*** |
| Observed features | **0.000666\*** |

**Supplemental Table S11.** Adonis test (PERMANOVA) to examine the effects of location on beta diversity of wild fly microbiome.

| Beta diversity metric | PR(>F) |
| --- | --- |
| Unweighted UniFrac distance | **0.001\*** |
| Bray-Curtis distance | 0.914 |
| Jaccard distance | 0.851 |
| Weighted UniFrac distance | **0.036\*** |

**Supplemental Table S12.** PERMDISP test to examine the effects of location on dispersion of wild fly microbiome.

| Beta diversity metric | PR(>F) |
| --- | --- |
| Unweighted UniFrac distance | 0.642 |
| Bray-Curtis distance | 0.628 |
| Jaccard distance | 0.51 |
| Weighted UniFrac distance | 0.626 |

**Supplemental Table S13:**
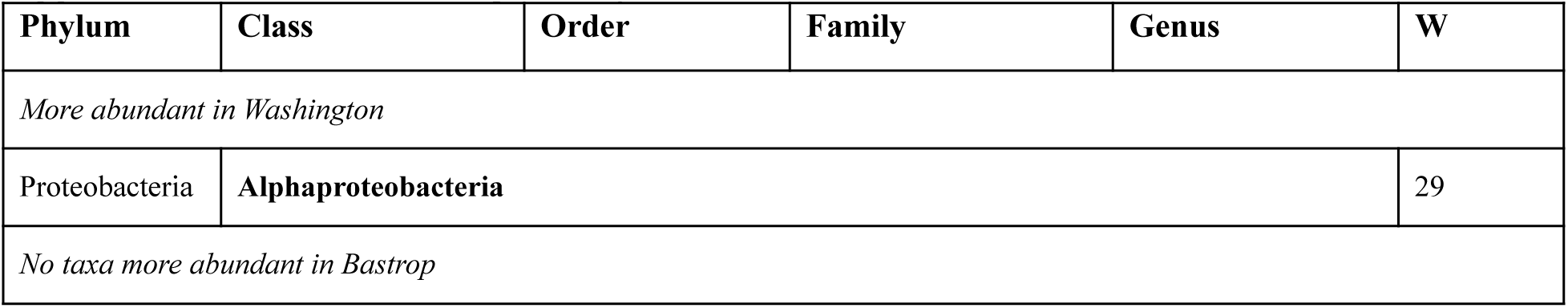
Taxa significantly differ in relative abundance across 2020 between two locations.

**Supplemental Table S14:** Taxa significantly different in relative abundance across 2021 between two locations.

| Phylum | Class | Order | Family | Genus | W |
| --- | --- | --- | --- | --- | --- |
| *No taxa more abundant in Washington* |  |  |  |  |  |
| *More abundant in Bastrop* |  |  |  |  |  |
| **Fusobacteriota** |  |  |  |  | 25 |
| **Desulfobacterota** |  |  |  |  | 22 |
| Firmicutes | **Clostridia** |  |  |  | 54 |
| Firmicutes | **Negativicutes** |  |  |  | 53 |
| Fusobacteriota | **Fusobacteriia** |  |  |  | 52 |
| Desulfobacterota | **Desulfovibrionia** |  |  |  | 52 |
| Bacteroidota | Bacteroidia | **Bacteroidales** |  |  | 138 |
| Firmicutes | Clostridia | **Oscillospirales** |  |  | 138 |
| Firmicutes | Clostridia | **Peptostreptococcales-Tissierellales** |  |  | 138 |
| Fusobacteriota | Fusobacteriia | **Fusobacteriales** |  |  | 137 |
| Firmicutes | Negativicutes | **Veillonellales-Selenomonadales** |  |  | 132 |
| Desulfobacterota | Desulfovibrionia | **Desulfovibrionales** |  |  | 129 |
| Firmicutes | Negativicutes | **Acidaminococcales** |  |  | 125 |
| Firmicutes | Clostridia | **Clostridia_UCG-014** |  |  | 120 |
| Firmicutes | Bacilli | **Erysipelotrichales** |  |  | 120 |
| Proteobacteria | Gammaproteobacteria | **Aeromonadales** |  |  | 120 |
| Firmicutes | Clostridia | **Clostridia_vadinBB60_group** |  |  | 116 |
| Verrumicrobiota | Lentisphaeria | **Victivallales** |  |  | 111 |
| Bacteroidota | Bacteroidia | Bacteroidales | **Bacteroidaceae** |  | 283 |
| Firmicutes | Clostridia | Oscillospirales | **Ruminococcaceae** |  | 273 |
| Bacteroidota | Bacteroidia | Bacteroidales | **Rikenellaceae** |  | 269 |
| Firmicutes | Bacilli | Lactobacillales | **Lactobacillaceae** |  | 266 |
| Fusobacteriota | Fusobacteriia | Fusobacteriales | **Fusobacteriaceae** |  | 264 |
| Bacteroidota | Bacteroidia | Bacteroidales | **Tannerellaceae** |  | 262 |
| Firmicutes | Clostridia | Peptostreptococcales<br>-Tissierellales | **Peptostreptococcaceae** |  | 257 |
| Bacteroidota | Bacteroidia | Bacteroidales | Bacteroidaceae | ***Bacteroides*** | 628 |
| Firmicutes | Bacilli | Lactobacillales | Lactobacillaceae | ***Lactobacillus*** | 615 |
| Firmicutes | Clostridia | Peptostreptococcales-Tissierellales | Peptostreptococcaceae | ***Romboutsia*** | 615 |
| Fusobacteriota | Fusobacteriia | Fusobacteriales | Fusobacteriaceae | ***Fusobacterium*** | 613 |
| Bacteroidota | Bacteroidia | Bacteroidales | Rikenellaceae | ***Rikenellaceae_RC9_gut_group*** | 612 |
| Proteobacteria | Gammaproteobacteria | Enterobacterales | Enterobacteriaceae | ***Escherichia-Shigella*** | 603 |
| Bacteroidota | Bacteroidia | Bacteroidales | Tannerellaceae | **‘-‘** | 593 |
| Firmicutes | Negativicutes | Weillonellales-Selenomonadales | Selenomonadaceae | ***Megamonas*** | 587 |
| Bacteroidota | Bacteroidia | Bacteroidales | Rikenellaceae | ***Alistipes*** | 572 |
| Bacteroidota | Bacteroidia | Bacteroidales | Prevotellaceae | ***Alloprevotella*** | 572 |
| Bacteroidota | Bacteroidia | Bacteroidales | - | **-** | 570 |

## Supplemental Figures

**Supplemental Figure S1:**
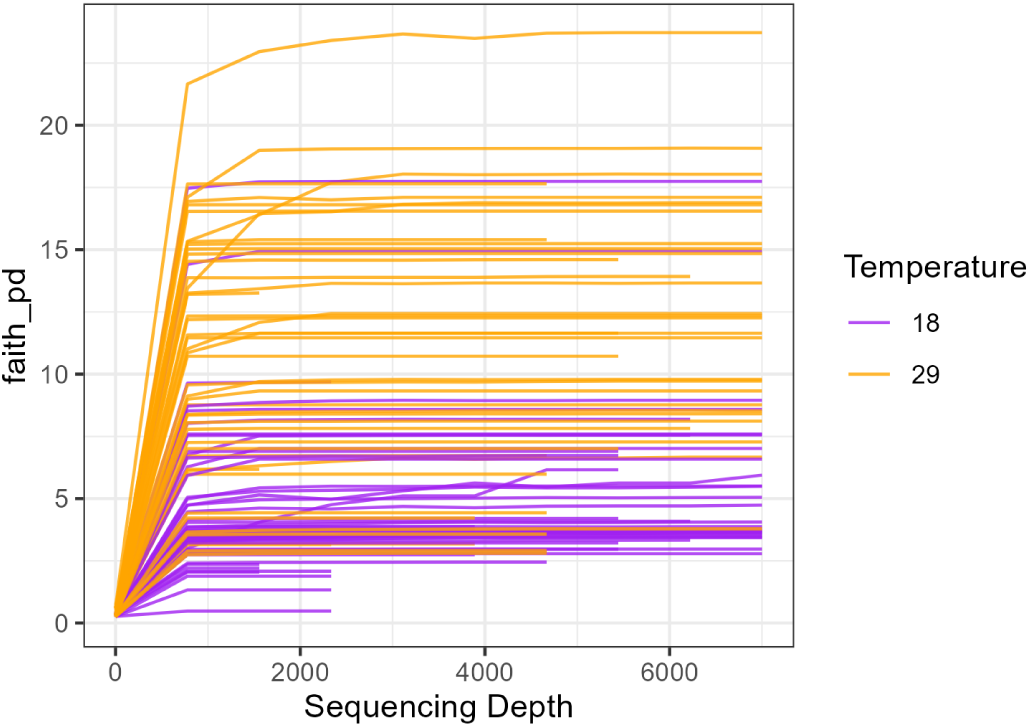
Rarefaction curve showing the individual lab-raised house fly. Each individual curve represents an individual house fly sample raised at either the 18°C (black) or 29°C (orange).

**Supplemental Figure S2:**
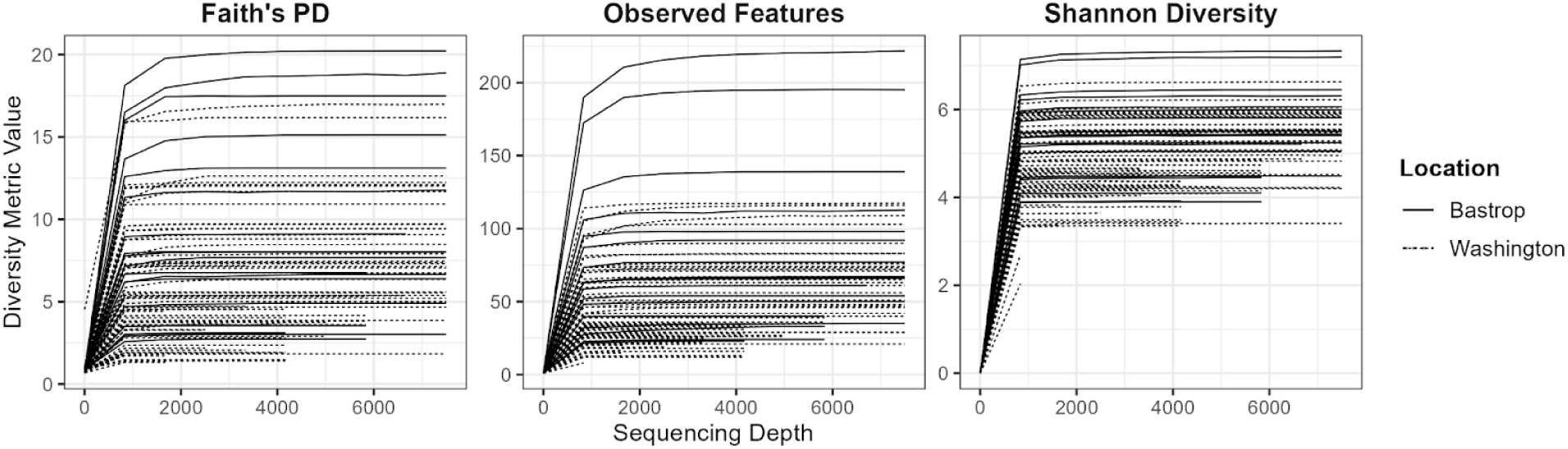
Rarefaction curve showing the individual wild-caught house fly. Each individual curve represents one house fly sample collected from either Bastrop (solid black) or Washington county (dashed black).

**Supplemental Figure S3:**
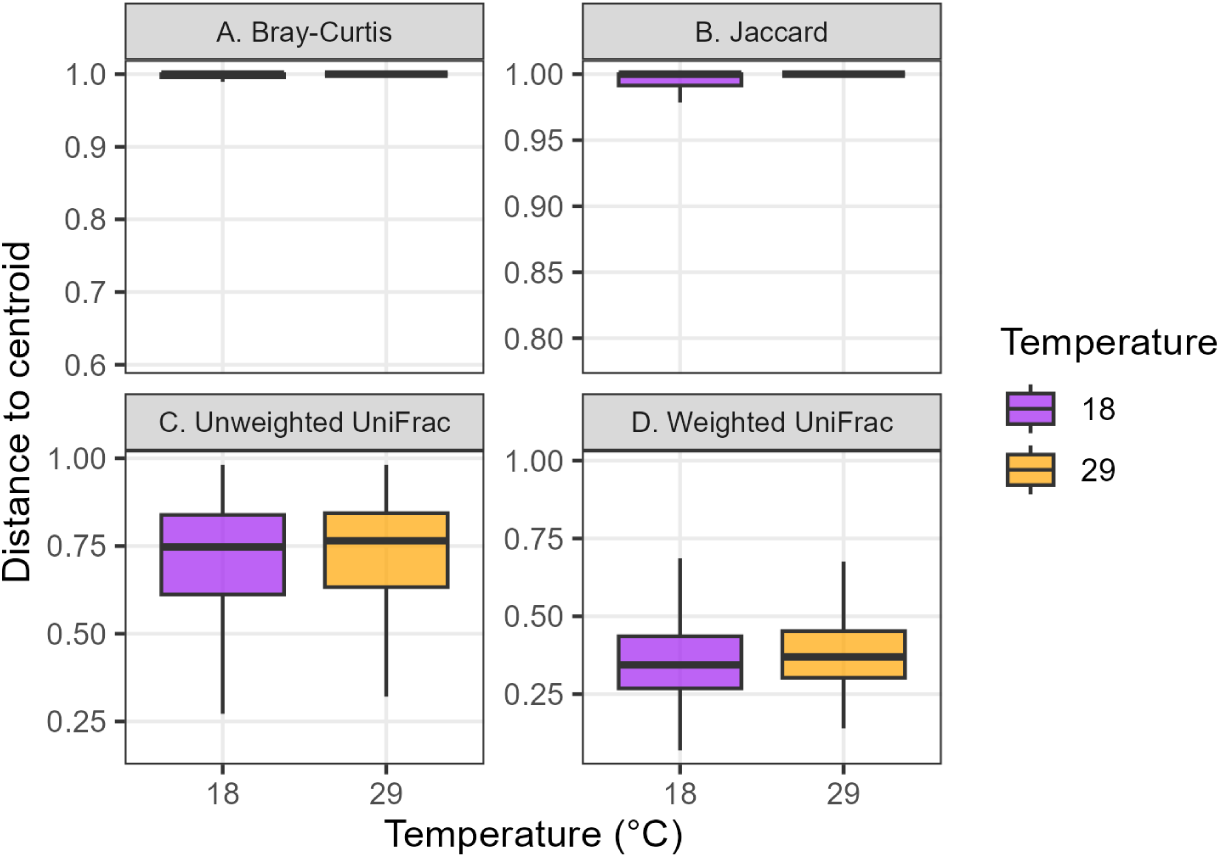
**Box plot showing dispersion between the lab-raised houseflies within each temperature for each diversity metric. The X and Y axes represent the temperature and distance to centroid respectively.**

**Supplemental Figure S4:**
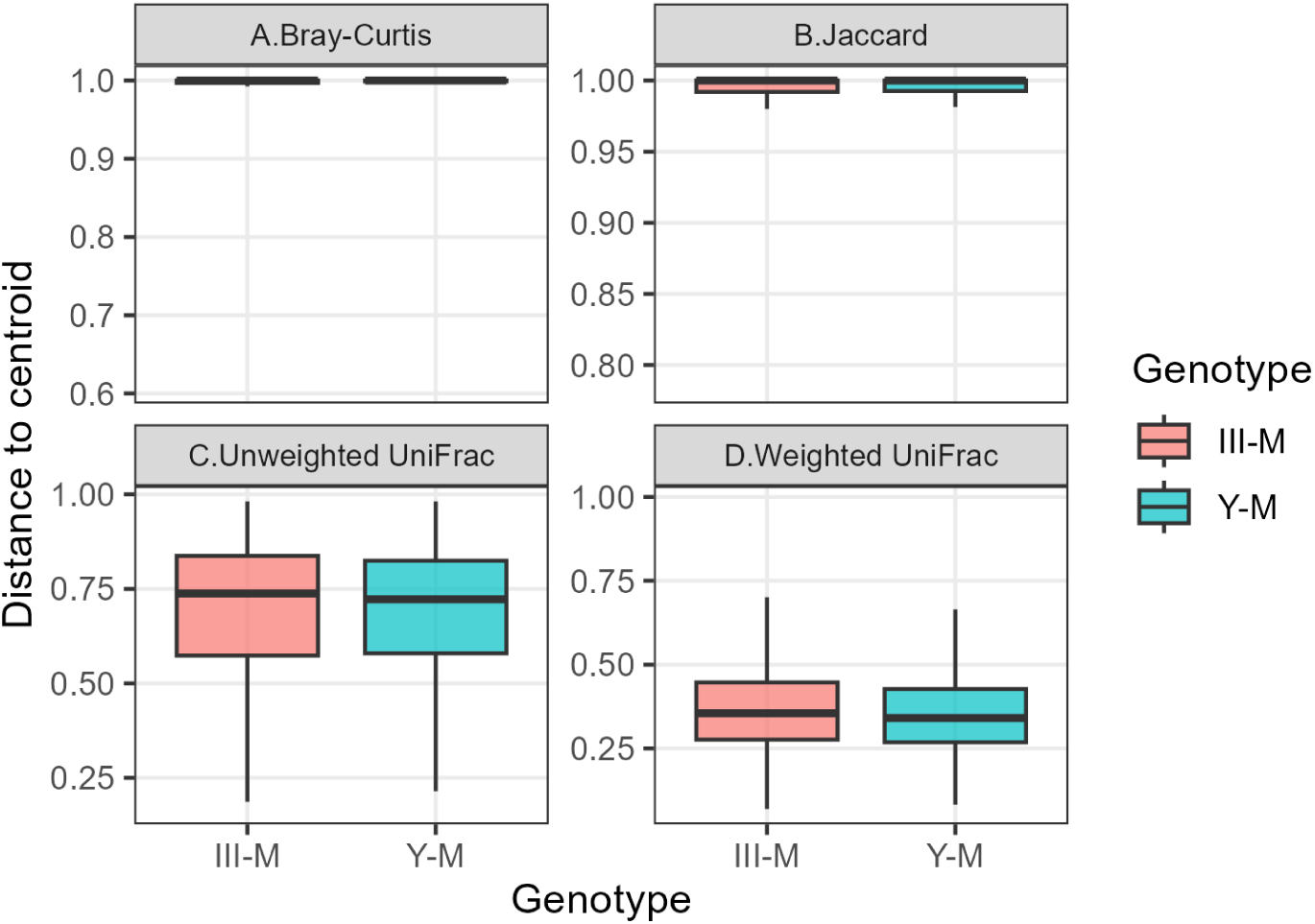
**Box plot showing dispersion between the lab-raised houseflies within each genotype for each diversity metric. The X and Y axes represent the temperature and distance to centroid respectively.**

**Supplemental Figure S5:**
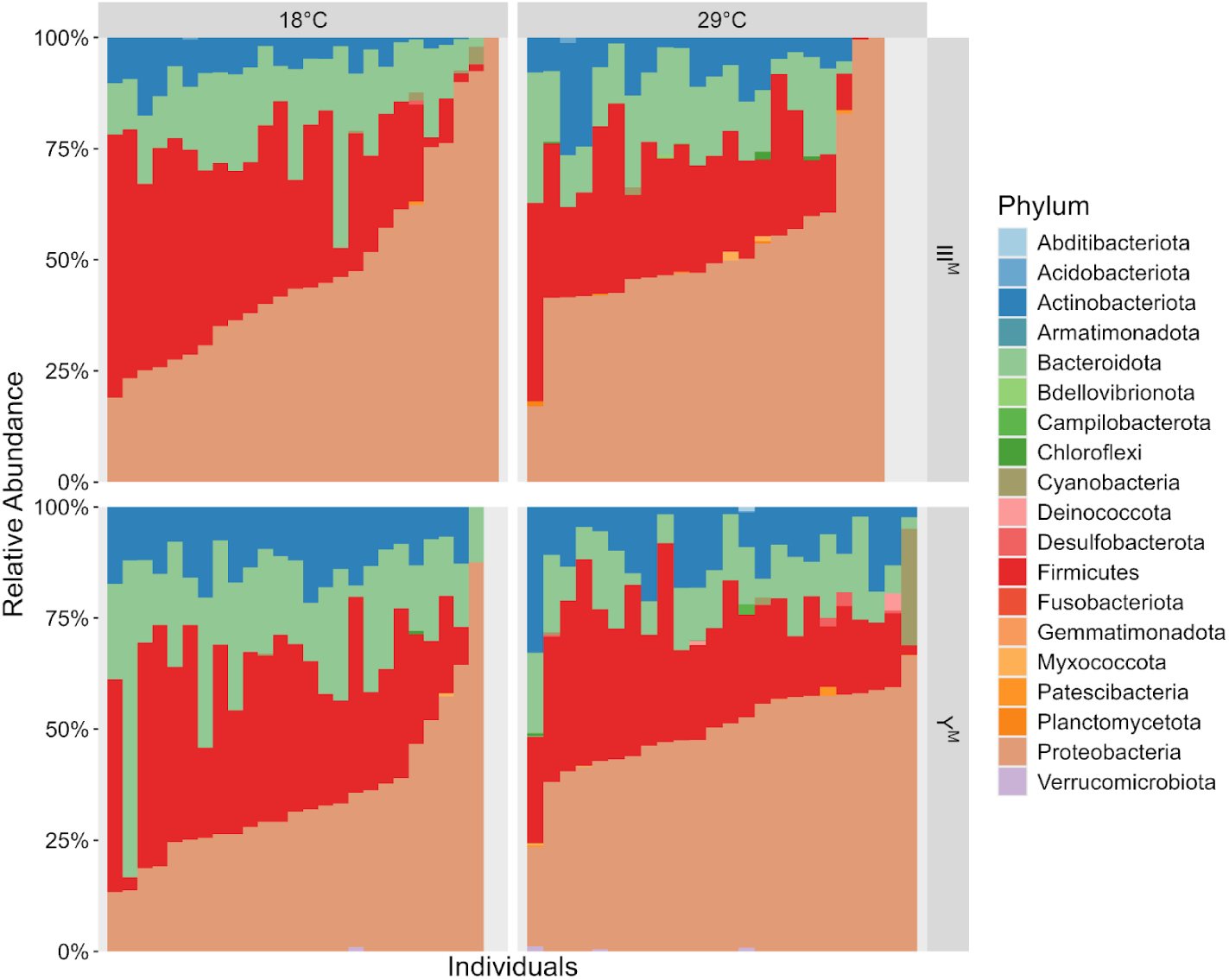
**Relative abundance of bacterial phyla in individual house fly samples, grouped by genotype and temperature. Bars show proportions of each phylum per sample, sorted by *Proteobacteria* abundance.**

**Supplemental Figure S6:**
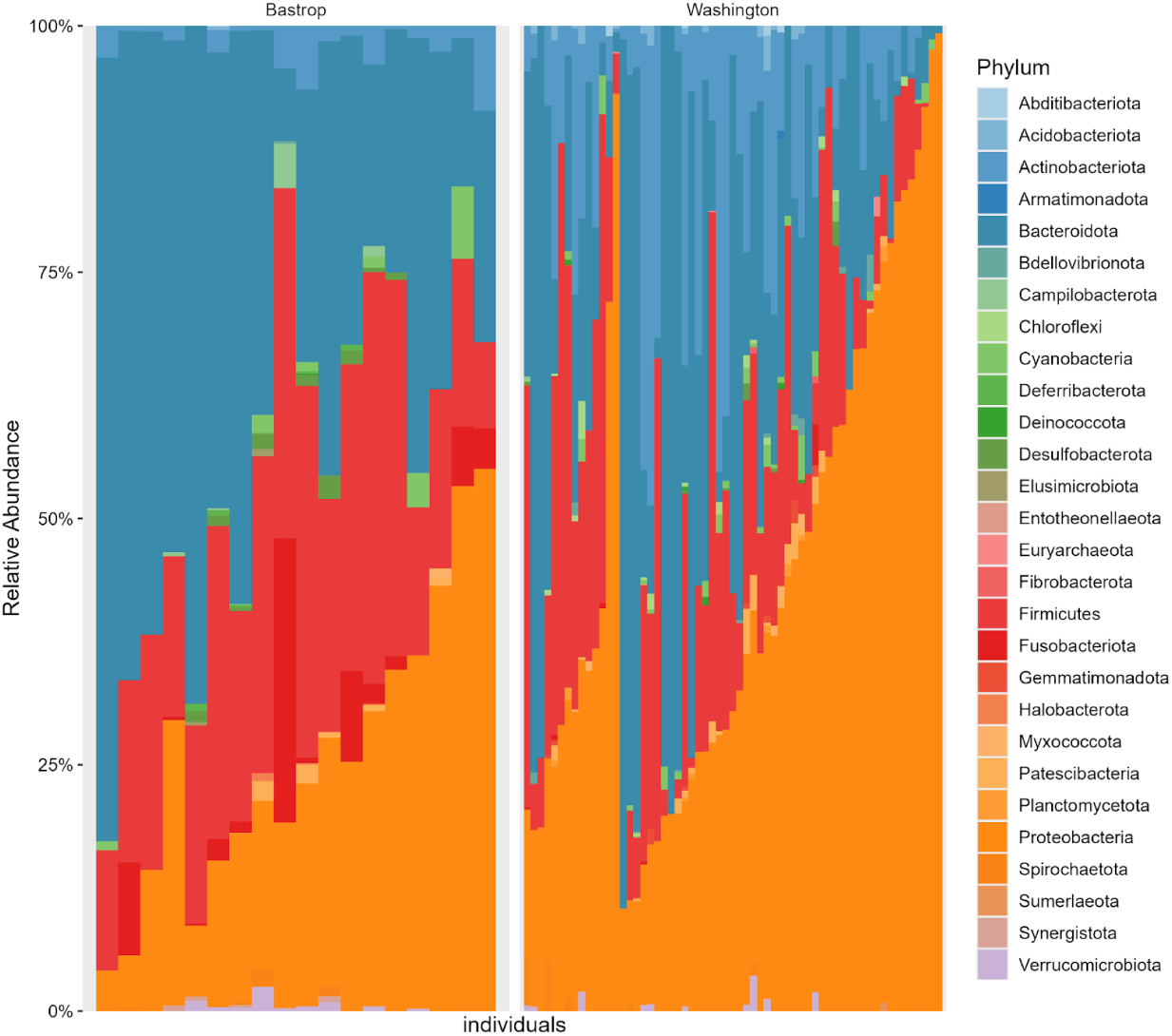
**Relative abundance of bacterial phyla of wild-caught flies from Bastrop and Washington Counties**.

